# The impact of respiratory gating on improving volume measurement of murine lung tumors in micro-CT imaging

**DOI:** 10.1101/823245

**Authors:** S. J. Blocker, M. D. Holbrook, Y. M. Mowery, D. C. Sullivan, C. T. Badea

## Abstract

Small animal imaging has become essential in evaluating new cancer therapies as they are translated from the preclinical to clinical domain. However, preclinical imaging faces unique challenges that emphasize the gap between mouse and man. One example is the difference in breathing patterns and breath-holding ability, which can dramatically affect tumor burden assessment in lung tissue. As part of a co-clinical trial studying immunotherapy and radiotherapy in sarcomas, we are using micro-CT of the lungs to detect and measure metastases as a metric of disease progression. To effectively utilize metastatic disease detection as a metric of progression, we have addressed the impact of respiratory gating during micro-CT acquisition on improving lung tumor detection and volume quantitation. Accuracy and precision of lung tumor measurements with and without respiratory gating were studied by performing experiments with *in vivo* images, simulations, and a pocket phantom. When performing test-retest studies *in vivo*, the variance in volume calculations was 5.9% in gated images and 15.8% in non-gated images, compared to 2.9% in post-mortem images. Sensitivity of detection was examined in images with simulated tumors, demonstrating that reliable sensitivity (true positive rate (TPR) ≥ 90%) was achievable down to 1.0 mm^3^ lesions with respiratory gating, but was limited to ≥ 8.0 mm^3^ in non-gated images. Finally, a clinically-inspired “pocket phantom” was used during *in vivo* mouse scanning to aid in refining and assessing the gating protocols. Application of respiratory gating techniques reduced variance of repeated volume measurements and significantly improved the accuracy of tumor volume quantitation *in vivo*.

## INTRODUCTION

Preclinical imaging has become an important tool for translating imaging techniques for the detection and monitoring of cancer. A growing application of small animal imaging is its utilization in supplementing a co-clinical trial (1, 2). However, there are critical challenges in the design and interpretation of translational animal studies that must be overcome to collect data which are meaningful in the context of clinical cancer research (3). These include, but are not limited to, subject size, imaging hardware, motion, etc.

We are developing quantitative imaging methods for use in the preclinical arm of a co-clinical trial investigating synergy between immunotherapy and radiotherapy in sarcoma (4). While magnetic resonance imaging is being employed to study primary lesion characteristics in the limbs of both patients and mice, a critical metric of therapeutic response in patients on study is metastasis-free survival determined through serial imaging of the lungs, which is the most common site for metastasis for this type of soft tissue sarcoma. To mimic the clinical study and patient standard of care, we are performing periodic micro-computed tomography (CT) scans of mouse lungs to detect and quantitate metastasis after treatment. Delineation of a significant trend in therapeutic response requires accurate and precise volumetric measurements of lung tumors, necessitating the implementation of respiratory gating techniques.

To understand the effect of respiratory gating on the sensitivity and precision of lesion detection and measurement, we performed a series of *in vivo* scans on primary lung tumor-bearing mice(5, 6). To better understand the effect of gating on true positive and false negative rates, we also used simulated tumor images, both gated and non-gated, which could be compared to a predetermined ground truth. Additionally, we evaluated the utility of retrospectively gated images as an alternative to prospective gating with reduced scan time. Finally, we have designed a clinically-inspired “pocket phantom” (i.e., a reference standard) for external use during scanning to aid in assessing gating protocols (7). We performed a series of *in vivo* scans on tumor-bearing mice while varying the animal’s position (test-retest), and performing the analyses in triplicate (repeated measures) to assess the effects of gating on tumor detection and volumetric assessment.

## METHODS

### Animal Model

Lung tumors were generated by intranasal administration of adenovirus expressing Cre recombinase (Adeno-Cre, Gene Transfer Vector Core, University of Iowa) into *LSL-Kras^G12D^; p53^FL/FL^* mice as described previously (5, 6). Mice with primary lung tumors were imaged at 12 weeks post Adeno-Cre infection, at which point multiple aggressive lung adenocarcinomas (∼0.5–1.5 mm in diameter) were present within each mouse. All animals were imaged at 24–30 weeks of age.

### Micro-CT

All micro-CT imaging was performed using a micro-CT system developed in-house (8, 9). Animals were scanned while breathing freely under anesthesia using 2–3% isoflurane delivered by nose-cone. A pneumatic pillow was positioned on the animal’s thorax and connected to a pressure transducer to monitor respiratory motion and inform gating. Body temperature was maintained with heat lamps and a feedback controller. The x-ray source parameters were 80 kVp and 40 mA with 10 ms exposures. A total of 360 views were acquired over a full 360° rotation. The reconstruction resulted in a 63 μm isotropic voxel size using the Feldkamp algorithm (9) followed by bilateral filtration to reduce noise (10). The micro-CT images were converted to Hounsfield units (HU) and saved in DICOM format. The radiation dose associated with a micro-CT scan was ~0.017 Gy per mouse, which is ~294 to 411 times less than the LD50/30 lethal dose (5-7 Gy) in mice (11).

### Phantom examinations for micro-CT

To assess image quality for our micro-CT scanner, we used a Micro-CT Phantom Kit that is commercially available (pureimaging.com). This kit includes multiple phantoms: i) a water phantom that is used for HU calibration as well as for measuring uniformity, noise and the noise power spectrum (NPS) assessed using volumes of interest measuring 64×64×64 voxels around the periphery and at three different Z positions; ii) a calcium hydroxyapatite (HA) phantom for bone density calibration that can also be used to verify CT number linearity; iii) a phantom containing 25 μm tungsten wires designed to assess the modulation transfer function (MTF) used for determining spatial resolution; iv) a multilayer phantom (also known as Defrise phantom) to assess the severity of cone beam artifacts when using approximate reconstruction algorithms.

### Prospective gating techniques

While clinical chest CT is performed during a single breath hold, projection data in micro-CT must be acquired over many breaths. Resulting motion can lead to image blurring which confounds measurement of small structures (such as developing metastases or primary lung tumors), making respiratory gating necessary to achieve reasonable accuracy. Respiratory gating can be performed prospectively or retrospectively (12). In prospective respiratory gating, a single respiratory phase (e.g., end expiration) can serve well to assess lung nodules. We were first to develop and implement combined cardiac and respiratory gating for micro-CT (13), which improves cardio-pulmonary imaging quality. However, the increased acquisition time which results from the addition of cardiac gating makes combined gating unfeasible for the large number of animals and frequent scan schedule required in the co-clinical trial. Consequently, we have used prospective gating to synchronize acquisition with respiration only, in which each projection is triggered when the respiratory signal breaches a user-defined threshold (Fig. 1). Thus, all projections are acquired in the same phase of the respiratory cycle (e.g., end inspiration), which minimizes motion artifacts and blurring in the reconstructions (13).

**Figure 1:**
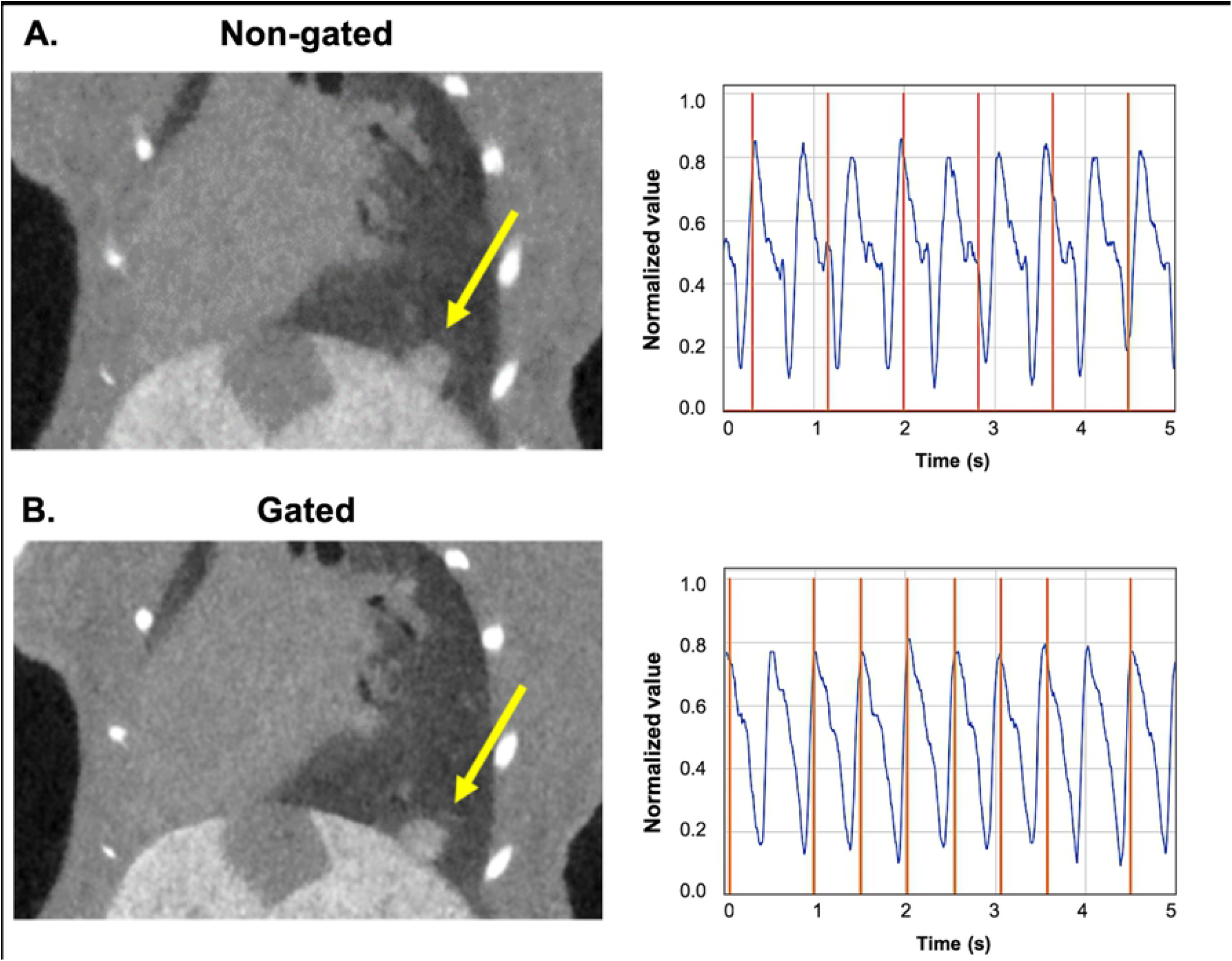
Prospective gating allows for synchronous image acquisition with breathing patterns. Micro-CT acquisitions without (A) and with respiratory gating (B). Shown are coronal reconstructed tomograms (left) presenting a lung lesion (yellow arrow). Breath rate (blue curve) and acquisition times (orange vertical lines) are shown for each gating technique (right). Note the sharper diaphragm and tumor margins in the gated acquisition.

### Simulated tumor nodules

We have examined the effect of respiratory gating on lung tumor detection and volumetric measurement using simulated nodules inserted into temporally-resolved micro-CT datasets. A C57BL/6 mouse without lung tumors was scanned four times to acquire four different prospectively-gated respiratory phases. Gated image volumes were reconstructed using Feldkamp’s algorithm (9) using 360 projections from each respiratory phase and processed with bilateral filtration (14). A lung mask was created for each respiratory phase by thresholding the image to remove bone and soft tissue. The lung mask was dilated and eroded to make the mask internally contiguous.

Three manual segmentations of real lung tumors were taken from previous micro-CT image sets for incorporation into the reconstructions (Fig. 2). To augment this small bank of tumors, they were subjected to transformations involving random rotations and skews in three dimensions, resulting in tumors with a range of sizes (0.03 to 0.28 mm^3^). Transforming the tumors altered their gray-values and textures. To address this, random high frequency Gaussian noise and Gaussian filtering were applied to the tumor masks to make the augmented tumors statistically like the originals.

**Figure 2:**
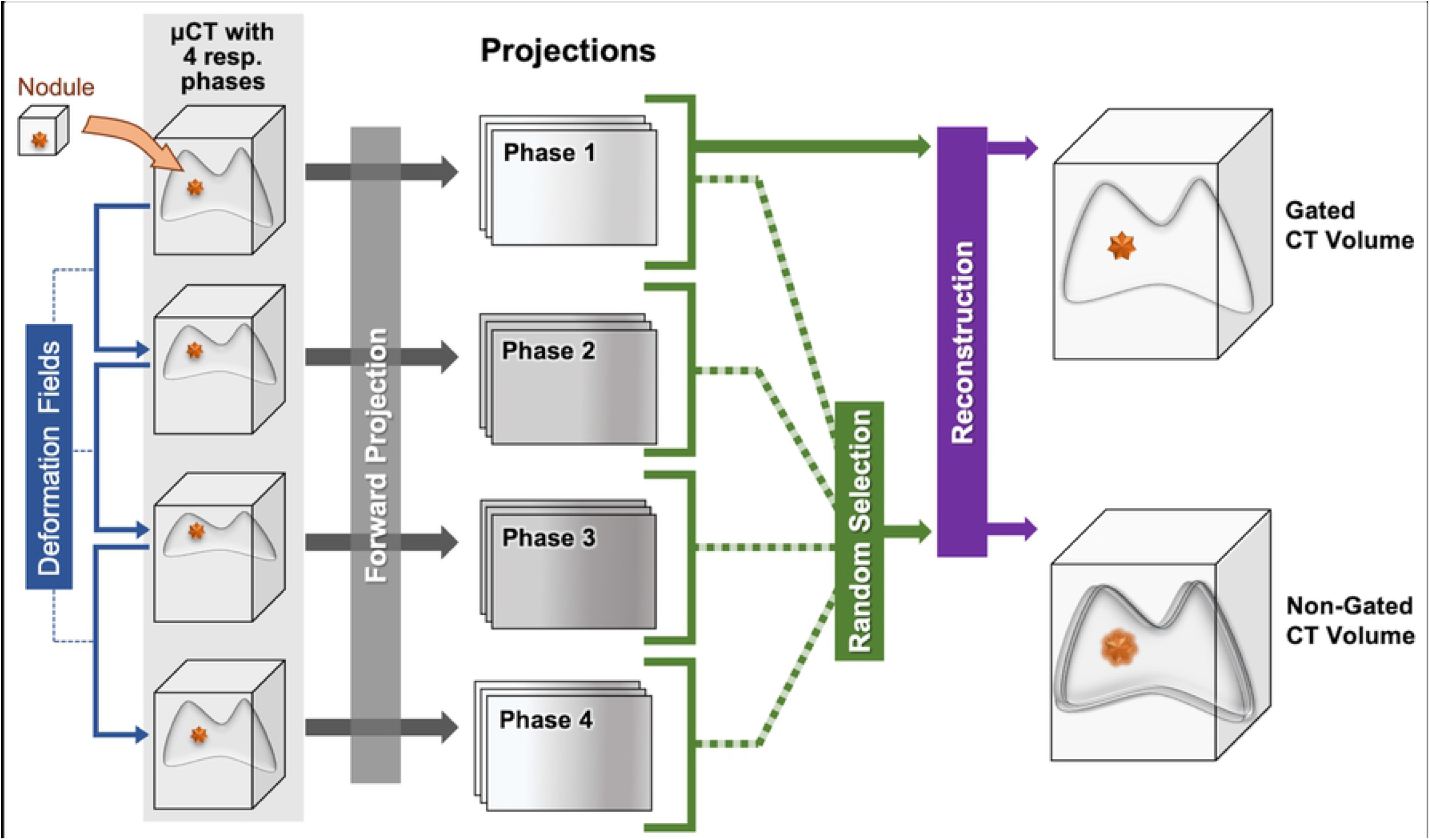
Process of adding simulated tumor to respiratory-gated and non-gated CT reconstructions. Deformation fields between respiratory phases are used to place tumors in volumes corresponding to different respiratory phases. Once placed, each volume associated with its lung phase is forward-projected in cone beam geometry. Random combinations of respiratory-gated projections are used to simulate non-gated acquisitions. The projections from a single lung phase create gated acquisitions.

The augmented tumors were randomly placed within the lung mask of the first respiratory phase. The permitted locations were such that the tumor did not have more than 10% overlap outside the lung-mask or with other tumors. In real data, lung tumors are often along the edges of the lungs and near other tumors. To simulate this property, tumor locations were weighted using the inverse of Euclidian distance to the nearest lung mask boundary (i.e., positions nearest an edge of the mask will be selected with higher frequency than those far from a mask boundary). To increase the likelihood of creating groups of tumors, previously placed tumor locations were removed from the lung mask, creating boundaries with high probability for additional tumors. Once all tumors were placed in the volume, the image was forward-projected in cone beam geometry to simulate CT sampling.

To simulate non-gated acquisitions, a deformation field was computed for the lungs for each reconstructed respiratory phase using the demons algorithm (15, 16). The deformation fields indicate how a tumor’s 3D position should change with respiratory motion. The magnitudes of the computed deformation fields within the lung were small due to difficulties for the algorithm to track low-contrast features in the lungs, however the direction vectors were consistent with observed data. To correct this, we applied a multiplicative factor to the deformation fields which provided a similar magnitude of motion as in the respiratory-gated acquisitions. Tumor placement in each respiratory phase was altered from the first respiratory phase by the modified deformation field. After tumor placement, each respiratory phase image was forward-projected in the scanner cone beam geometry. Non-gated reconstruction was simulated by randomly selecting combinations of projections from each of the four respiratory phases. For gated simulations, only the first respiratory phase was used for reconstruction, mimicking the product of active breath gated acquisitions *in vivo*. Each image set contained gated and non-gated reconstructions with twelve simulated tumors.

### Retrospectively gated image generation and analysis

Images of five tumor-bearing mice were acquired with prospective respiratory gating and without (non-gated). Respiration was monitored throughout the non-gated acquisition, and the resulting projections were divisible into four breath phases. Projections from a single respiratory phase (phase 0) were used as the retrospectively gated CT images for tumor segmentation. Unfortunately, retrospective gating produces an irregular angular distribution of projections per phase; thus, when using Feldkamp’s reconstruction algorithm, streaking artifacts affect the image quality of the tomographic reconstructions (17).

In a retrospective gating strategy, the projection images corresponding to a chosen respiratory phase are also fewer in number. Thus, an iterative reconstruction is needed to reduce the artifacts associated with missing angles. Reconstructions performed with fewer projections generally appear noisier than well-sampled reconstructions. To improve the reconstruction of retrospectively-gated data, we have used a sophisticated iterative reconstruction algorithm based on the split Bregman method with the addresidual-back strategy and with 4D bilateral filtration for regularization (18). In essence, we solve the problem of temporal reconstruction (i.e., four different respiratory phases) by minimizing the following penalized, weighted least squares objective function:

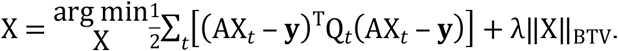

Each column of the reconstructed data, X, represents the reconstructed volume at a different respiratory phase, indexed by *t*. The columns of X are related to the acquired projection data, **y**, through the CT system projection matrix, A, and a time-point-specific, diagonal weighting matrix, Q_*t*_. The weights on the diagonal of Q_*t*_ are repeated for each detector pixel within a single projection. The regularization term is controlled via the regularization parameter λ. For retrospective gating, projection weights are assigned based on Gaussian basis functions which are centered on the respiratory phase to be reconstructed and which have a full-width-at-half-maximum equal to the integration time of the projection data, scaled to span the complete respiratory cycle. The iterative reconstruction was initialized with temporally-weighted least-squares evaluated for each time point, *t*, (n_t=4_ respiratory phases) and with the biconjugate gradient stabilized convex solver (BiCGSTAB (19)). For regularization we minimize bilateral total variation (BTV) through the application of bilateral filtration (BF, (20)). The filtration domain is expanded to include the respiratory phase point proceeding (t-1) and following (t+1) the time point being filtered.

The signal-to-noise (SNR) was measured in the prospectively gated, retrospectively gated, and non-gated images via regions of interest (ROIs) drawn in the liver and empty space. Three tumors from each mouse were selected in the prospectively gated image, varying the size and relative location as much as possible. Tumor isocenter coordinates were recorded for consistent localization across segmentations. Measurements were performed in triplicate in each image type, alternating between prospective, retrospective, and non-gated. Sets within each triplicate were performed with at least one day in between measurements, to avoid serial-segmentation biases in edge detection. Resulting tumor volumes were compared within each individual image (test-retest of measurements to determine measurement variance), as well as between image types (to determine accuracy and reproducibility under different gating conditions).

### Phantom studies

Previously, we have used x-ray radiopaque beads surgically glued to the abdominal diaphragmatic surface and cardiac ventricles of anesthetized, mechanically ventilated rats to describe the range of motion using digital microradiography with respiratory gating methods (21). For this study, a mouse phantom similar to the CT human “pocket phantom” (5) was fabricated using a number of polyethylene spheres (http://www.cospheric.com/) with diameters of ~0.7 mm. The spheres were embedded in a packaging foam, which imitates the appearance of lung parenchyma, and loaded into a microcentrifuge tube. The complete phantom was positioned on the animal chest to freely move with respiration, and analysis of the beads was used as a reference to quantify respiratory gating performance (Fig. 3).

**Figure 3:**
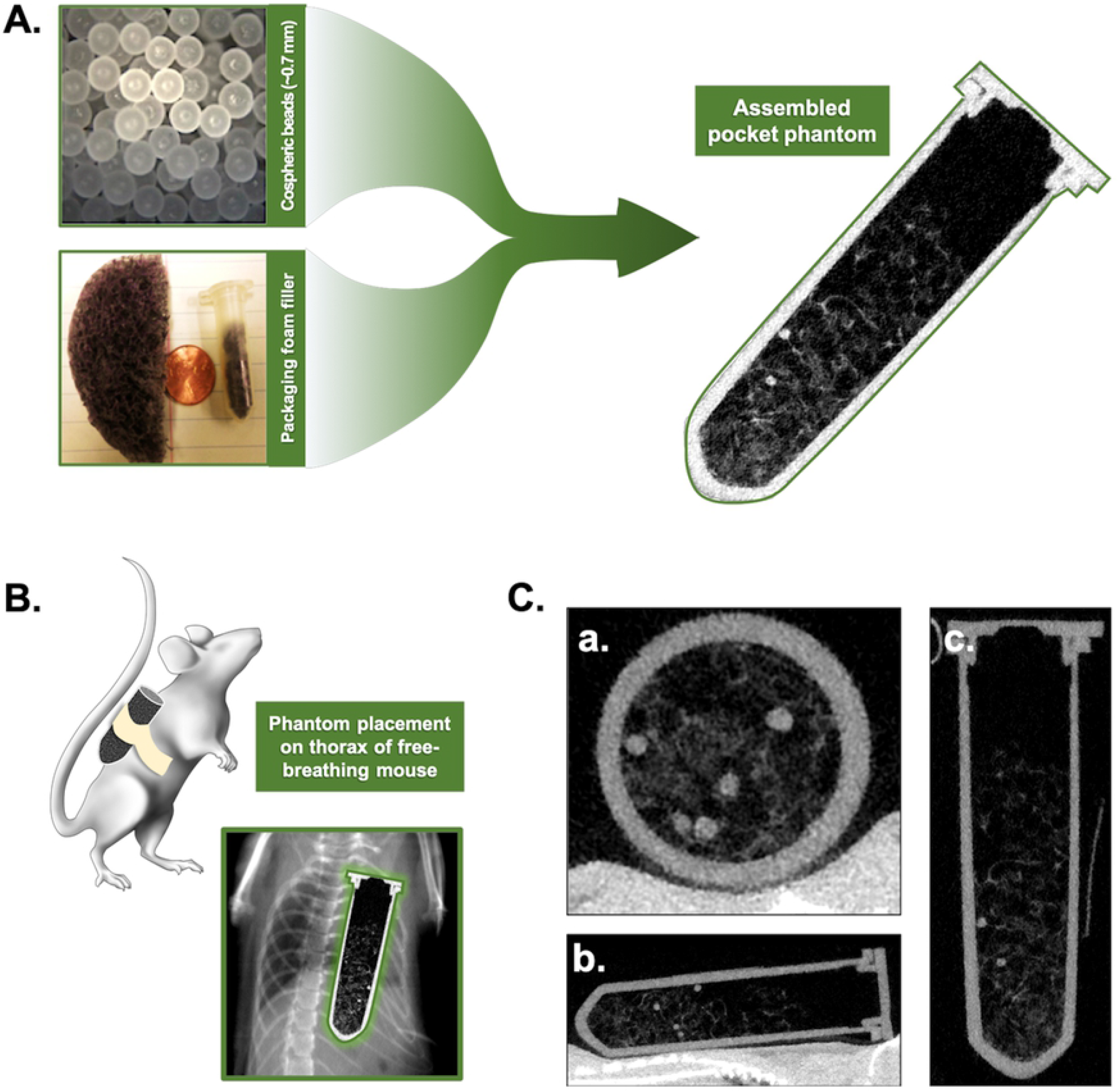
Design of a pocket phantom for evaluation of respiratory gating. The phantom was constructed using packaging foam to mimic the lung parenchyma in which a number of polyethylene spheres were inserted (A). The constructed pocket phantom was fixed to the thorax of free-breathing mice, allowing for uninhibited motion throughout the respiratory cycle (B). Examples of the micro-CT images of the pocket phantom (C) are shown in the axial (a.), coronal (b.), and sagittal (c.) view.

### Mouse scans

For preliminary *in vivo* lung tumor studies, we scanned the same mouse in three adjusted positions (repositioned on scan bed), acquiring images with and without gating in each position. A non-gated scan requires about approximately 3 minutes and a prospectively gated scan takes about 5 minutes to acquire. Post-mortem *ex vivo* scans of the same mouse were also performed in triplicate and used as the standard for defining tumor volumes and segmentation precision.

Subsequent in vivo imaging was performed by acquiring one gated and one non-gated image (as described above) in a single position.

### Image analysis

All measurements, including segmentations and SNR measurements, were performed in triplicate. A collection of identifiable tumors *in vivo*, phantom pellets, or simulated tumors were identified in each of the scan sets. Volumes were calculated via manual segmentation in OsiriX DICOM viewer or 3D Slicer (https://www.slicer.org/) image analysis software. For simulated images, the reader was blinded and asked to identify as many tumors as possible, as well as to segment each lesion to calculate a volume. Subsequently, the reader performed manual segmentations on the binary label map containing all simulated tumors (“truth”) to measure the precision of the segmentation method. Each tumor identified in the gated and non-gated images (including true positives and false positives) was compared to the binary label map to calculate detection sensitivity and precision, as well as volumetric accuracy. *In vivo*, tumors and phantom pellets were identified for each condition (gating, no gating, post-mortem) and each repositioning on the cradle, resulting in nine measurements per volume of interest.

### Establishment of a Quantitative Imaging Biomarkers Alliance (QIBA)-inspired profile for volumetry

In order to understand the effect of precision on day-to-day tumor assessments for our co-clinical trial, we have implemented a QIBA-inspired profile for mouse lung tumor volumetry in CT with gating techniques based on tumor size (22, 23). In a pilot group of tumor-bearing mice, images were acquired with and without prospective respiratory gating. After scanning, the animals were sacrificed and post-mortem images were acquired. Lung nodules (n=13) were identified and measured in triplicate in each of the 3 conditions. Coefficients of variance (CV) were calculated for each measured tumor volume in gated, non-gated, and post-mortem conditions. The tumors were then divided into 3 groups based on size (as measured post-mortem). Within-subject coefficients of variance (wCV) were determined for each size group in gated and non-gated images (24), and used to calculate the 95% confidence intervals (CI) as follows:

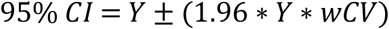

where Y is the measured tumor volume, and wCV is dependent upon the size of Y.

To assess the utility of the calculated 95% CIs, an independent group of tumor-bearing animals were scanned with and without respiratory gating. Three tumors of varying sizes were identified in each mouse (n=15), and volume measurements were repeated three times for each tumor/condition. The results were plotted against the calculated 95% CIs to determine if the size-dependent prediction of precision reasonably reflected variance of repeated measures across a range of tumor sizes.

### Statistical analysis

Analysis of variance (ANOVA) calculations were performed to examine the effect of gating on volume reproducibility. Bland-Altman analyses were used when comparing acquisition techniques in simulated and retrospectively gated images. Statistical significance was defined as: *p<0.05.

## RESULTS

Technical performance of the micro-CT scanner was assessed with phantom scans (Fig. 4). The mean noise values (standard deviation) in the water phantom were around 68 HU (Fig.4 A). Note that the levels of noise are much higher than in clinical CT due to the higher resolution. However, these noise levels are acceptable when imaging a high contrast organ such the lungs and using very low radiation dose as is required for longitudinal imaging. The line profile shows good uniformity with no cupping. The system has linear response (Fig. 4 B) and spatial resolution given by the MTF at 10% is 5.5 line pairs/mm (Fig.4 C). The NPS measured in the water phantom indicates some structured noise. The noise power decreases significantly at high spatial frequencies (Fig. 4 D). We also show that with our scanner, the cone beam artifacts are acceptable when using Feldkamp algorithm with a circular scanning trajectory (Fig. 4 E). Our cone beam geometry creates cone angle of ~10 degrees.

**Fig. 4:**
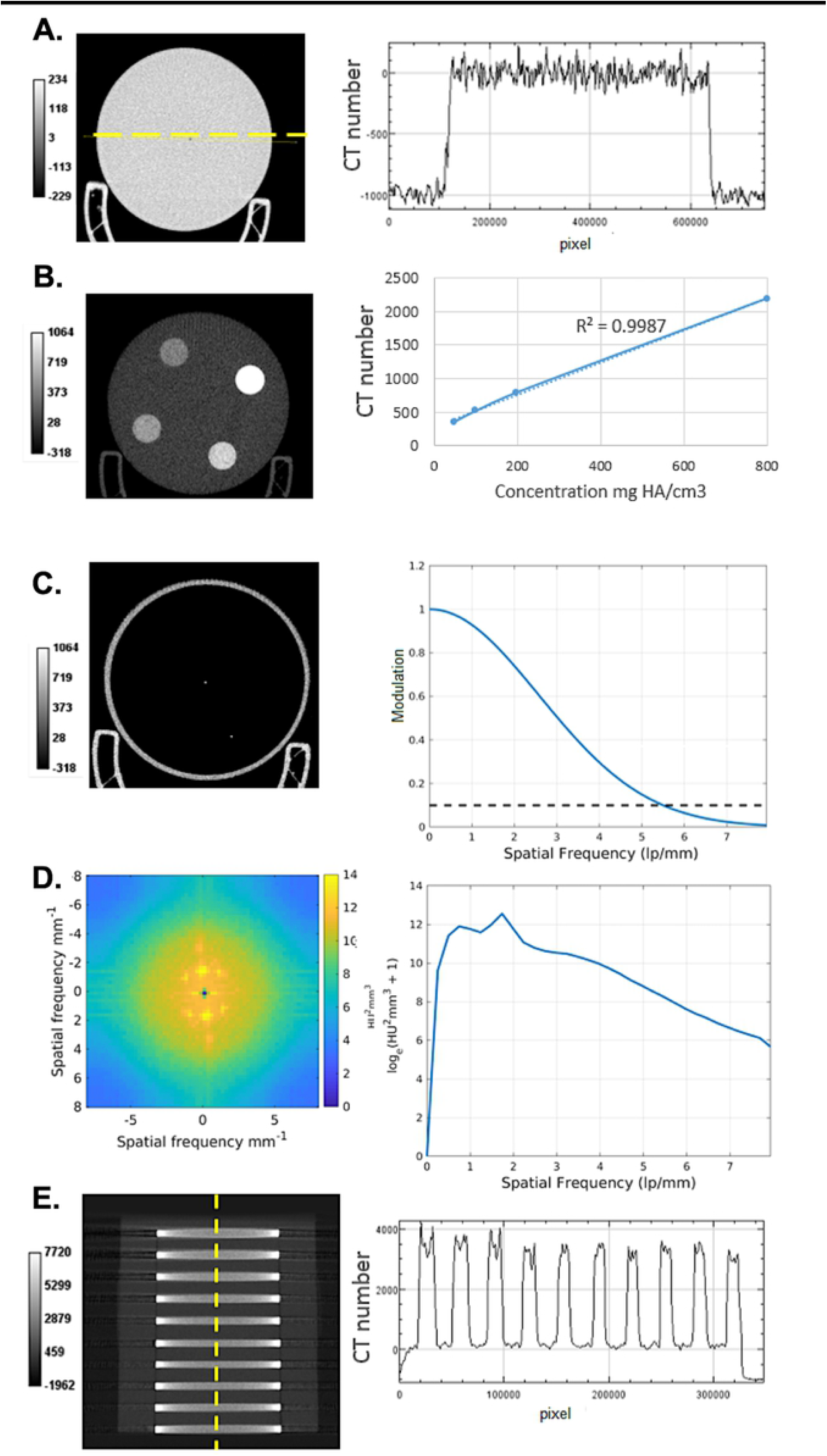
Phantom assessment of the micro-CT scanner performance. Shown is a slice and a line profile through the water phantom (A). The CT linearity was obtained by scanning a phantom containing 4 inserts with various concentrations of calcium hydroxyapatite (B). The spatial resolution was assessed with a wire phantom (C). Shown also is the image and the radial profile of NPS obtained in the water phantom (D). Finally, a coronal slice and a line profile of the multi-layer Defrise phantom is shown (E). No substantial cone beam artifacts are present.

### Respiratory gating reduces variance and improves accuracy of tumor volume measurements *in vivo*

To quantify the impact of respiratory gating on tumor volume measurements, selected lung lesions in mice with primary tumors were segmented in triplicate from images acquired with gating, without gating, and after sacrifice (motion-free control). Tumors of varying size which were locatable in all three acquisition conditions were selected for analysis (Fig. 5). The mean volume calculated in non-gated images was significantly different (*p<0.05, calculated via ANOVA) from the post-mortem values in 2 out of 4 tumors (Fig. 5 B-C). Gated volumes did not differ significantly, suggesting that gating improves accuracy of volume measurements compared to non-gated images. In all cases, the mean volume calculated from the non-gated image was greater than both the post-mortem and gated images. On average, the variance in volume calculations was 2.9% in post-mortem images, 5.9% in gated images, and 15.8% in non-gated images (Fig. 5 D). Thus, prospective gating reduces variance compared to non-gated images in repeated measures, demonstrating improved volume measurement precision. Gating did not result in higher variance or lower accuracy than non-gated images when calculating tumor volumes in any case. Taken together, these data suggest that in free-breathing mice, prospective gating reduces motion-based blurring and improves tumor volumetry precision and accuracy.

**Fig. 5.**
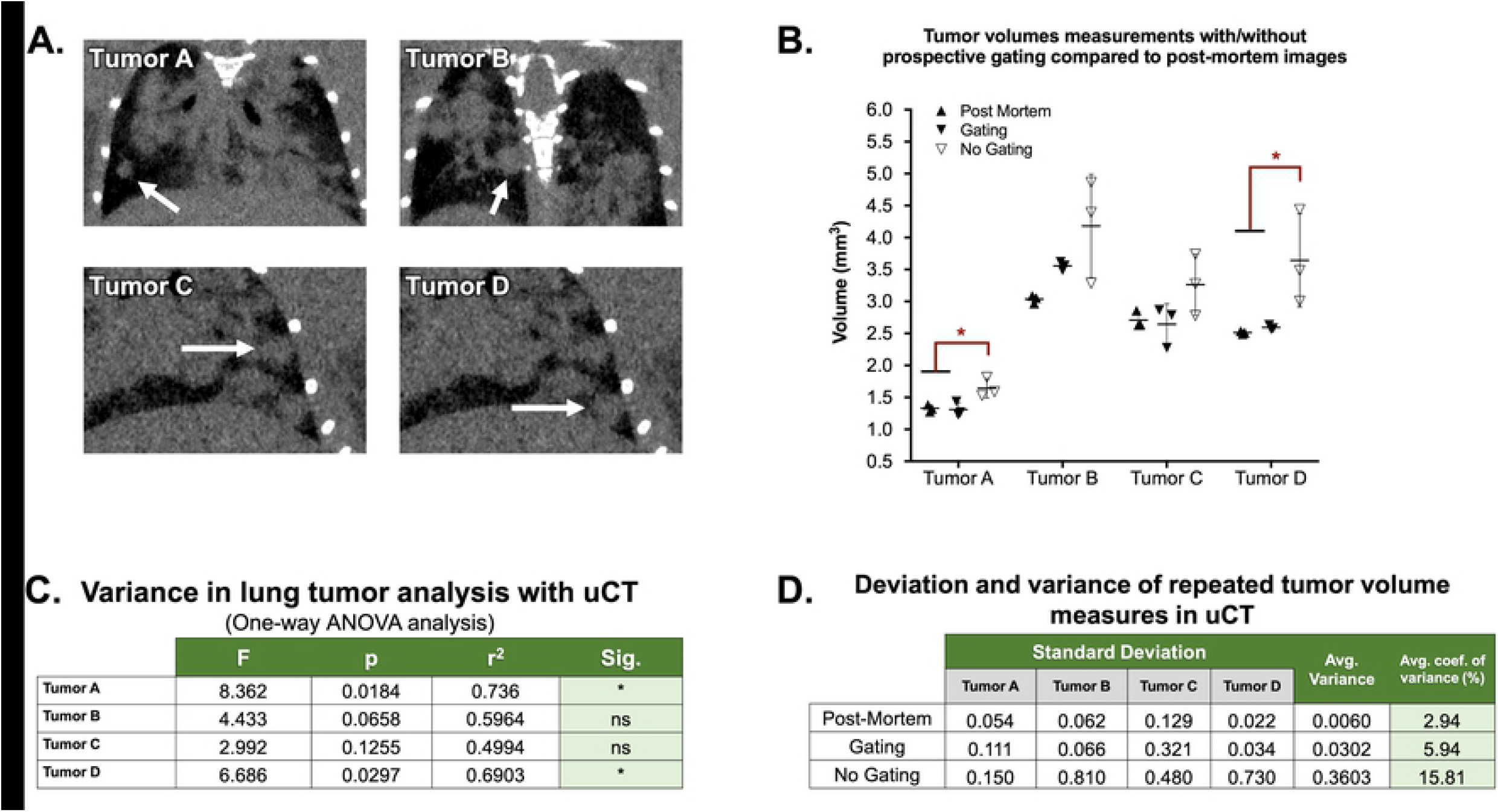
Respiratory gating improved the accuracy and precision of tumor volume measurements in the lung. Volumes were calculated for a collection of tumors (A) in 3 positions per condition, including a non-gated image, a gated-image, and an image acquired post-mortem (standard). When accounting for variance, volumes from non-gated images were significantly different from post-mortem standards in half of cases, while gated image volumes did not differ significantly (B; *P<0.05, ANOVA). The resulting ANOVA table describes the deviation and difference in means between acquisition condition per tumor (C). Coefficients of variation were calculated for each condition and averaged for each acquisition type (D).

From data acquired in animals scanned before and after sacrifice, we constructed a QIBA-inspired profile for lung tumor segmentation in CT images acquired with and without gating on our scanner. Given the dependency of segmentation precision or accuracy on nodule size, the predicted confidence intervals were calculated for ranges of tumor volumes observed on study. As expected, wCV and confidence intervals were generally reduced for volume assessments in gated images compared to non-gated images (Fig. 6 A). This disparity increased with rising nodule volume, where the precision of gated volumetry improved while non-gated precision remained unchanged. To assess the utility of the derived confidence intervals for gated and non-gated volumetry, we performed test-retest triplicate segmentations of multiple tumors in five mice. In gated and non-gated images, the calculated 95% confidence intervals were successful in predicting variance of volume measurements based on tumor size, with only one notable exception in a non-gated segmentation of a large tumor (Fig. 6 B). Thus, the derived 95% confidence intervals provide a reasonable metric for estimating the probable variance in tumor volume measurements acquired with our lung CT protocols. This is of immense value in a large-scale, co-clinical trial setting, where high animal numbers make persistent measures of precision infeasible. In addition, the need for longitudinal assessments requires a means of understanding the differences in precision as a tumor changes size, without the validation of post-mortem ground truth.

**Figure 6.**
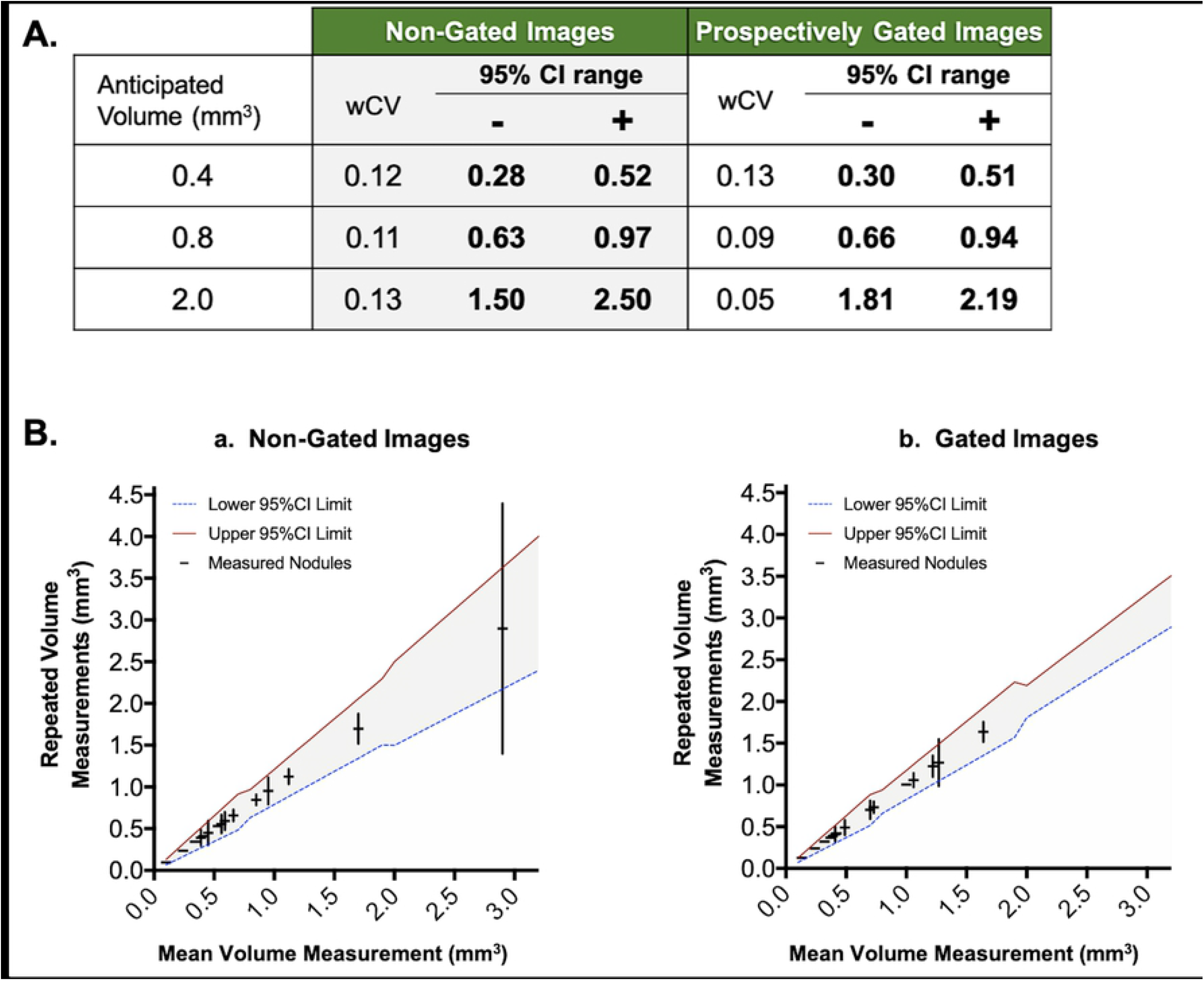
QIBA-inspired profile for predicting tumor volumetry precision across variable tumor sizes in gated and non-gated images. Shown is a table of sample anticipated tumor volumes for which the wCV and upper/lower 95% CIs was determined in both non-gated and gated acquisitions (A). Triplicate tumor measurements (plotted as mean ± standard deviation) from an independent cohort of mice were plotted against the calculated 95% CIs (red and blue bounding lines) (B), demonstrating the validity of the size-dependent confidence prediction for both non-gated (a) and gated (b) images.

### Simulated lung nodules clarify the sensitivity and specificity of tumor detection in micro-CT of mouse lungs with and without respiratory gating

To assess false negative rates and limits of detection, we utilized our lung micro-CT datasets containing simulated tumors which can be compared to a binary label map, or absolute truth (Fig. 7). Simulated tumor volumes were measured in triplicate on the original binary label map (to quantitate any variance due to segmentation drawing), the non-gated images, and the gated images (Fig. 8 A). Precision of repeated volume measurements was frequently superior in the gated acquisitions compared to the non-gated measurements. Additionally, the accuracy of tumor volume calculations in gated images was consistently better than in non-gated images, with multiple non-gated volume calculations deviating significantly from the binary label map (truth).

**Figure 7.**
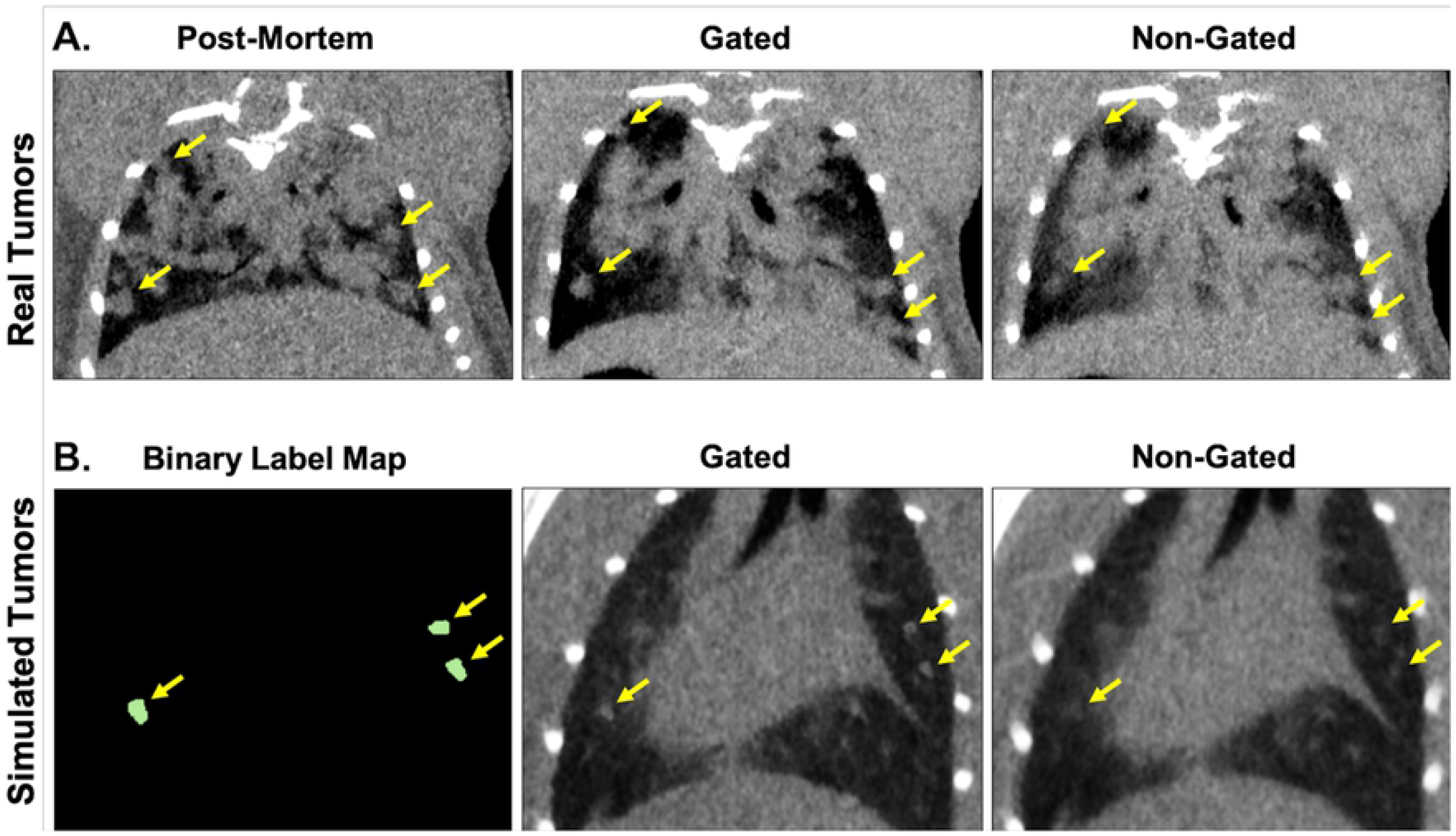
Demonstration of simulated lung tumor images compared to in vivo images of tumor-bearing lungs. *In vivo* lung images from a tumor-bearing mouse demonstrate the difference between non-gated and gated images compared to post-mortem tumor detection (A). Simulated tumors are created from segmentations of real tumors, which are projected into the lungs of a healthy, free-breathing mouse (B). Simulated tumors mimic the appearance and location of tumors seen within *in vivo* images, as well as the effects of respiratory gating on tumor edge definition, but have the advantage of a binary label map which serves as “truth”. Real and simulated tumors are indicated with yellow arrows on *in vivo* and simulated acquisitions, respectively.

**Figure 8.**
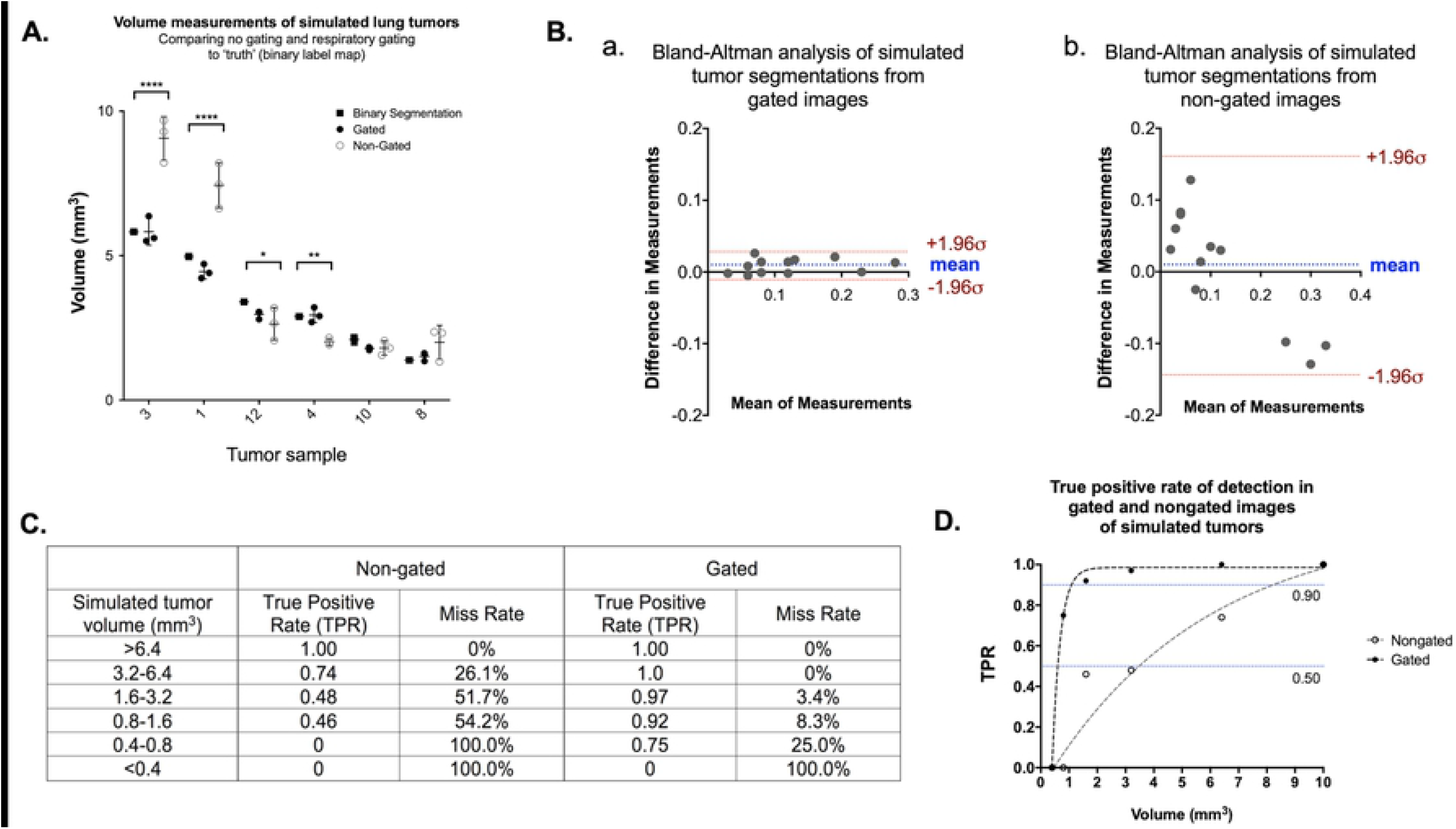
Simulated tumors in CT of free-breathing mouse lungs demonstrate enhanced sensitivity, precision, and volume accuracy when identifying tumors in gated images compared to non-gated images. Variably sized samples of measured simulated lung lesions are shown, including triplicate measures of binary label maps (truth), gated images, and non-gated images (A; ANOVA, *p<0.05). Bland-Altman plots of gated (B.a.) and non-gated (B.b.) tumor segmentations suggest agreement with segmentations of the binary label map, with a size-dependent bias noted in non-gated image segmentations. True positive rate (TPR; sensitivity), Miss rate (False negative rate), Positive predictive value (precision), False discovery rate (False positive rate), and F1 (harmonic mean of precision and accuracy) are shown for gated and non-gated images of simulated tumors of varied sizes (C). TPR as a function of simulated lesion size, with TPRs of 0.9 and 0.5 indicated (blue lines) (D).

Bland-Altman analysis comparing volume measurements from binary label maps with those derived from gated and non-gated acquisitions suggested that both acquisition types resulted in volume measurements of reasonable agreement with the ground truth (Fig. 8 B). However, the trend seen in the plot of non-gated acquisitions, demonstrated by a linear negative slope, suggests that volume calculations from these images are subject to a tumor size-dependent bias. Specifically in non-gated images, large simulated tumors and small simulated tumors were frequently over- and under-estimated, respectively.

Finally, with the availability of ground truth we were able to identify the limitations in sensitivity of both gated and non-gated acquisitions. The occurrence of both false negative and false positive lesion identification was greater in images without respiratory gating. The true positive rate (TPR; sensitivity) in correctly identifying tumors was improved with the application of gating to 0.97, compared to 0.56 in non-gated images (Fig. 8 C). Similarly, the positive predictive value (PPV; precision) of gated images was 1.0, compared to 0.71 in non-gated images. When comparing limits of detection of TPR = 0.9 (the size of lesion at which TPR drops below 0.9), gating maintained reliable detection down to tumor volumes of approximately 1.0 mm^3^, while non-gated images were not reliable in positive prediction at volumes below approximately 8.0 mm^3^ (Fig. 8 D).

### Retrospective gating techniques improve volume measurement accuracy compared to non-gated images

Datasets including prospectively gated and non-gated images were acquired in five mice. Retrospective images were generated from the non-gated images by selecting projections from a single phase of respiration. First, signal-to-noise (SNR) measurements were taken by selecting ROIs in the liver and empty air surrounding the mouse (Fig. 9 A). Prospectively gated and non-gated images demonstrated similar SNRs, with prospective gating being slightly better. Retrospectively gated images showed SNR of approximately half of prospectively gated images (Fig. 9 B). This was expected, given that retrospectively gated images although reconstructed using an iterative approach use mostly one fourth of the data, resulting in more noise. Nevertheless, we note that iterative reconstruction does help compared to filtered back projection for the same amount of data.

**Figure 9.**
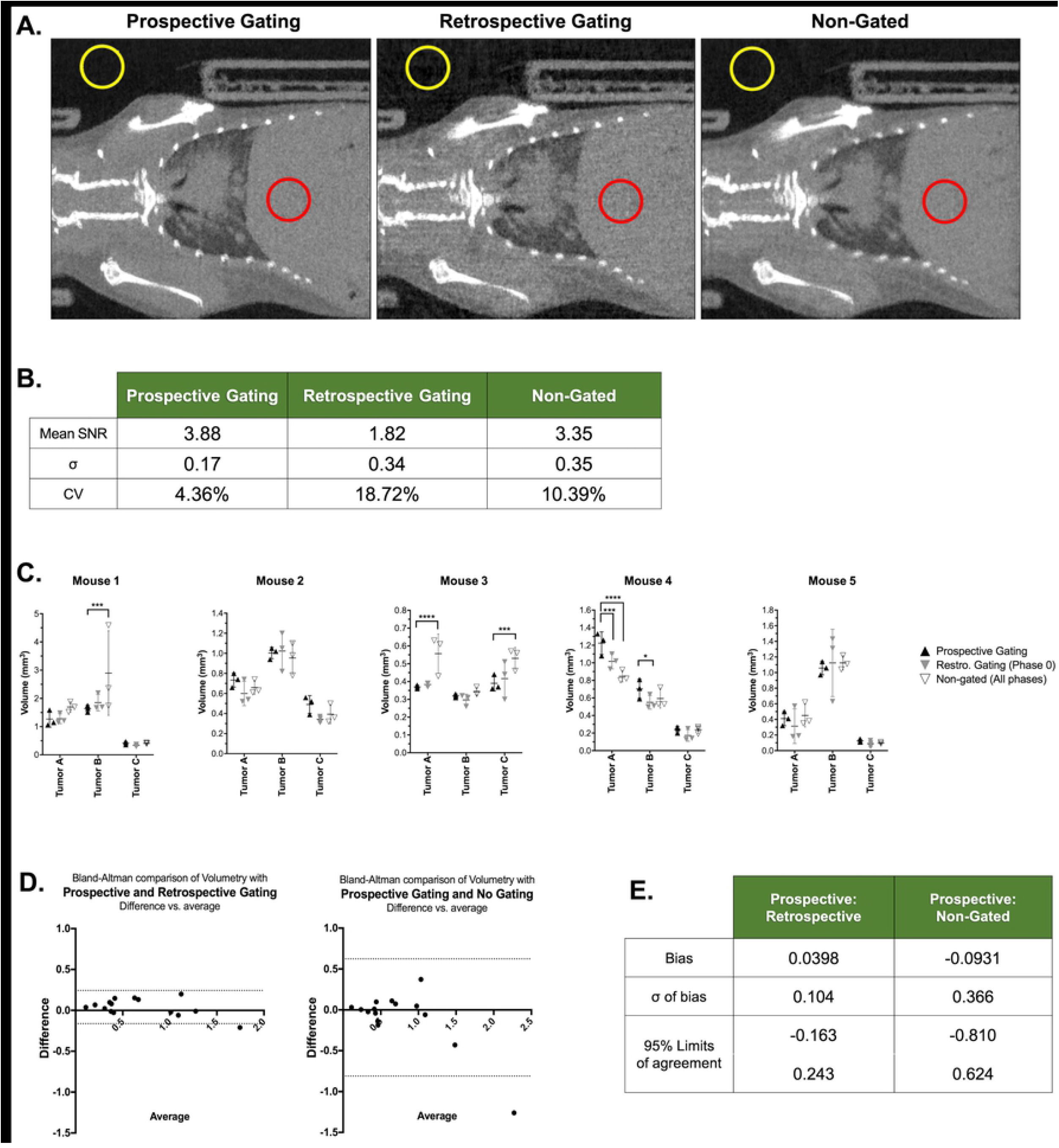
Retrospective gating improves tumor volume accuracy compared to non-gated images. Prospectively gated, retrospectively gated, and non-gated images were acquired in five mice bearing lung tumor lesions (A). SNR measurements were calculated from liver (red) and empty space (yellow) ROIs to determine mean SNR and variance (B). Three tumors in each mouse were selected and segmented in triplicate under each scan condition, and results were compared with ANOVA (C). Bland-Altman comparisons to prospective gating were performed for retrospective gated and non-gated tumor analyses (D & E).

Three tumors from each mouse were measured in triplicate in all three gating conditions, with each measurement occurring at least one day apart to avoid bias. Based on ANOVA comparisons, there were four tumors in which the non-gated image resulted in significantly different volume measurements compared to prospectively gated images (Fig. 9 C). Retrospectively gated images differed significantly from prospective gating in the case of two tumor volume measurements, one of which was also mis-measured in non-gated images. Bland-Altman comparison of both alternatives to prospective gating demonstrated that retrospective gating resulted in better agreement, with a smaller bias and reduced limits of agreement (Fig. 9 D-E). Overall, retrospective gating improved volume accuracy relative to non-gated images, but was not a perfect substitution for prospective gating. This is likely due to the variance in measurements caused by a lower SNR. Taken together, these data suggest that retrospective gating helps alleviate some, but not all, of the motion-related issues with tumor volumetry with a shorter acquisition time than prospectively gated lung CT.

### Use of a pocket phantom demonstrates respiratory motion in real time

Pellets contained in a pocket phantom on the thorax of tumor-bearing mice were used to assess the effect of prospective respiratory gating in real time during *in vivo* imaging. The variance of pellet volume measurements was determined from three repetitions in each of the following conditions: no gating, respiratory gating, post-mortem (motion-free) (Fig. 10). Three identifiable pellets were selected to mimic conditions that may be encountered during lung tumor measurement, including a solitary pellet, a pellet adjacent to the tube wall, and a pellet directly adjacent to another pellet.

**Figure 10:**
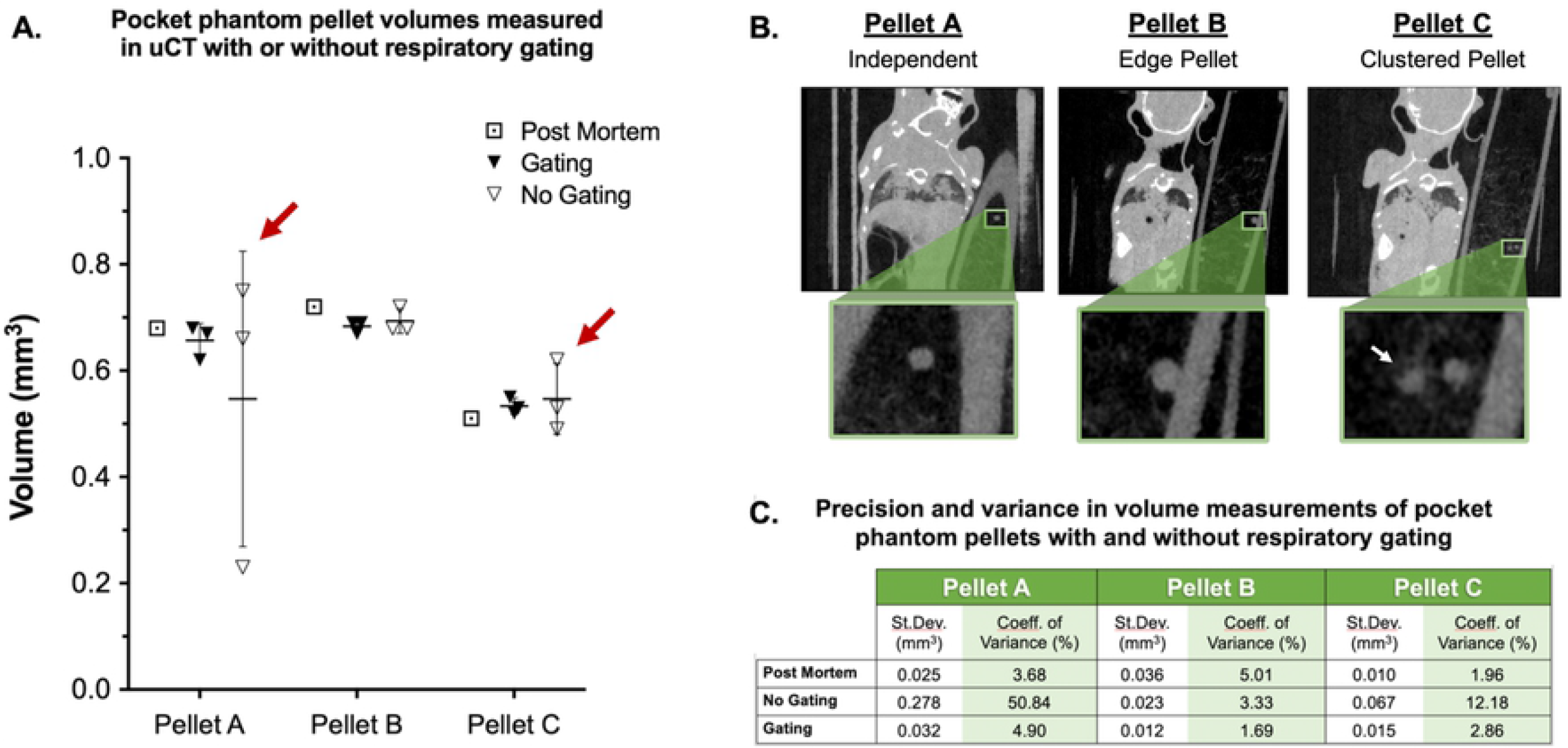
The effect of gating on volumetric output and measurement variance. Three pellets were measured three times in each acquisition condition, and volumes were calculated and compared to post-mortem measurements (A). Pellets were selected to mimic conditions regularly encountered during lung tumor analyses (B). Variance among measured volumes was greatest in non-gated images (red arrows in A), while gating resulted in precision similar to postmortem measurements (C).

## DISCUSSION AND CONCLUSIONS

The goal of this work was to assess the performance and utility of respiratory-gated CT for detection and measurement of tumors in mouse lungs to serve a co-clinical imaging study. When compared to clinical CT, the first challenge in implementing preclinical imaging is ensuring adequate and comparable instrumentation. The performance of the micro-CT scanner was found to be adequate for imaging of small tumor nodules in the lungs of mice. We note that there are important constraints on radiation dose levels which are intentionally very low to minimize any influence of longitudinal scanning on the outcome of the experiments.

Unlike clinical chest CT, in which patients may hold their breath to reduce motion during acquisition, breathing motion in micro-CT of mouse lungs is a persistent barrier to small lesion detection. The development of lung metastases is a critical metric for assessing therapeutic response in our co-clinical trial of sarcoma, making the ability to discern differences in small lesions between treatment arms paramount. Thus, we employed respiratory gating techniques to lessen the effect of motion in the lungs, increase visibility of small lesions, and improve tumor volume measurement. *In vivo* studies demonstrated that tumor segmentations performed in images with prospective respiratory gating more accurately measured the volume of tumors than non-gated segmentations, compared to post-mortem controls. In multiple instances, non-gated tumor volumes were significantly different than control measurements, while gated volumes never differed significantly. Further, gated segmentations were consistently more precise than those measured in non-gated images. These data support the use of prospective respiratory gating to improve tumor analysis in the lungs of free-breathing mice.

While post-mortem tumor measurement can serve as a motion-free control, the volumetric limits of detection with or without gating remain undefined. Specifically, postmortem controls do not provide a complete assessment of false negative and false positive rates, even though motion is not present. To delineate false negative rates, and how gating techniques affect them, we utilized images which contained simulated tumors that visually mimicked real tumors, projected into healthy, free-breathing micro-CT images of the mouse lung. Most importantly, the size, shape, and location of simulated tumors which appear in the final projections are known, providing a definitive ground truth to which analyses can be compared. Using this system, we were able to interrogate the sensitivity and precision of tumor detection in our micro-CT images, as well as quantitate the advantage of respiratory gating techniques with respect to lesion size. In fact, respiratory gating improved overall sensitivity and specificity of tumor detection compared to non-gated projections. Further, accuracy of tumor volume measurement was consistently better in gated images, while non-gated images demonstrated size-dependent discrepancies in volumetric accuracy.

While prospective respiratory gating significantly improved tumor detection and measurement accuracy/precision, there is a time cost for acquiring prospectively-gated images. Thus, we evaluated retrospective gating of non-gated images as an alternative to prospective gating. Conceptually, retrospective gating would reduce motion-related artifacts and blurring, improving lesion detection/measurement with the same short acquisition time of a non-gated image. We found that, indeed, retrospective gating did improve accuracy of tumor volumetry compared to non-gated images. However, retrospectively-gated images were consistently noisier, resulting in less precise measurements than both non-gated and prospectively-gated images. Thus, the utility of retrospective gating for tumor volume measurement lies somewhere between acquisitions without gating and prospectively gating. It is also worth noting that while some time is saved during acquisition compared to prospective gating, long reconstruction times (> 30 mins) associated with iterative reconstruction algorithms must be considered. Still, retrospective gating could serve as a reasonable alternative to prospective gating for those hoping to improve non-gated lung imaging without sacrificing scan time. We note that with more sophisticated reconstruction iterative algorithms, the performance of the retrospectively gated images can be improved (25).

Finally, given the impact of successful gating on image quality, we sought to create and implement a “pocket phantom” with which we could assess gating success on a scan-to-scan basis. We measured beads contained within the pocket phantom, which moved with the mouse as it breathed, with the same methods used to measure tumors. As was seen with lung lesions, gating improved precision when repeatedly measuring the volume of multiple pellets. The use of these pocket phantoms represents a simple and elegant technique by which users can monitor gating success *in vivo*. Specifically, these phantoms may be used as internal controls to identify instances of inadequate gating, such as in the case of weak or inconsistent respiratory signal.

Overall, we have demonstrated methods for respiratory gating for improving tumor detection and measurement accuracy in the lungs of free-breathing mice. Further, we have provided rigorous assessment of preclinical gating techniques, highlighting the advantages and shortcomings of multiple methods for correcting lung motion in the context of a co-clinical cancer imaging study.

## ACKNOWLEDGEMENTS

All work was performed at the Duke Center for In Vivo Microscopy supported by the NIH National Cancer Institute (R01 CA196667, U24 CA220245). We would like to acknowledge Drs. David Kirsch for providing the mice for the animal experiments and Yi Qi for helping with the animal imaging.

